# AI-MHC: an allele-integrated deep learning framework for improving Class I & Class II HLA-binding predictions

**DOI:** 10.1101/318881

**Authors:** John-William Sidhom, Drew Pardoll, Alexander Baras

## Abstract

**Motivation:** The immune system has potential to present a wide variety of peptides to itself as a means of surveillance for pathogenic invaders. This means of surveillances allows the immune system to detect peptides derives from bacterial, viral, and even oncologic sources. However, given the breadth of the epitope repertoire, in order to study immune responses to these epitopes, investigators have relied on *in-silico* prediction algorithms to help narrow down the list of candidate epitopes, and current methods still have much in the way of improvement.

**Results:** We present Allele-Integrated MHC (AI-MHC), a deep learning architecture with improved performance over the current state-of-the-art algorithms in human Class I and Class II MHC binding prediction. Our architecture utilizes a convolutional neural network that improves prediction accuracy by 1) allowing one neural network to be trained on all peptides for all alleles of a given class of MHC molecules by making the allele an input to the net and 2) introducing a global max pooling operation with an optimized kernel size that allows the architecture to achieve translational invariance in MHC-peptide binding analysis, making it suitable for sequence analytics where a frame of interest needs to be learned in a longer, variable length sequence. We assess AI-MHC against internal independent test sets and compare against all algorithms in the IEDB automated server benchmarks, demonstrating our algorithm achieves state-of-the-art for both Class I and Class II prediction.

**Availability and Implementation:** AI-MHC can be used via web interface at baras.pathology.jhu.edu/AI-MHC

## Introduction

The ability for T-cells to recognize various epitopes is of paramount importance to mounting a potent immune response and ultimately protecting the host (Cell, 1994). The relevance for understanding the ‘epitome’ for humans to viruses, bacteria, and even various cancers has been vital for advances in vaccine development, understanding how pathogens escape immune recognition, and even predicting how cancer patients will respond to immunotherapy(Ott *et al.,* 2017; Timm *et al.,* 2004; Łuksza *et al.,* 2017). Despite how much is known about epitope production including processing by the immunoproteasome, transport into the endoplasmic reticulum (ER), and binding and presentation via major histocompatibility (MHC) molecules, prediction of presented epitopes to the immune system is still a difficult task(Vyas *et al.*, 2008; Neefjes *et al.*, 2011).

The complexity of the task has led many groups to use advanced methods in machine learning and artificial intelligence to learn patterns in known MHC-binding peptides in order to recognize these patterns when seen in unknown peptides(Nielsen *et al.*, 2003; Andreatta and Bioinformatics, 2015). Artificial neural networks (ANN’s) have been employed by some of the leading algorithms to date to act as feature extractors in order to recognize patterns(Nielsen *et al.*, 2008; Andreatta *et al.*, 2015). Artificial neural networks, due to their flexibility in terms of changing their capacity, serve as universal function approximators, and therefore can learn patterns difficult for humans to pick up on. Building on the principal of using neural networks, groups have recently begun to utilize a type of neural network architecture termed convolutional neural networks (CNN’s) which were originally developed for the purpose of image classification where features in an image can be found in different locations and different orientations. By being translationally invariant to features, these networks put together the presence of multiple features in an image in order to make a decision as to what object is present in the image(LeCun *et al.*, 2015). This concept as applied to sequence analysis has been exploited in analyzing DNA-protein binding domains as well as predicting HLA Class I binding(Zeng et al., 2016; Vang and Bioinformatics, 2017; Han and bioinformatics, 2017).

While these most recent advances in neural network architectures have improved the accuracy of these algorithms, there are still areas for improvement. As a general shortcoming, most neural-network based methods of conducting MHC-binding predictions create several models across different alleles and different sequence lengths. The result of this process is that while the entire data set of known allele/peptide pairings is large, the data becomes split between models where each model can only learn sequence features for a subset of the peptides. However, it is known that neural networks, especially deep learning models, show the most increase in performance when more data is provided for training. *Andreatta et. al* demonstrated that using a gapped-sequence alignment method, they could feed variable length sequences into a fixed-input neural network by providing an additional parameter that specified the length of the original sequence (L ≤ 8, L = 9, L = 10, L ≥ 11), showing an improvement in MHC Class I binding prediction from being able to leverage more data in one model.

In order to best leverage the amount of data available for known MHC binding, we developed Allele-Integrated MHC (AI-MHC), a unified architecture capable of predicting binding for either all Class I or all Class II alleles, regardless of sequence length. By allowing MHC allele to be an input into the network and joining this with a global max pooling operation following convolutions across the peptide sequence, our architecture is able to leverage the most amount of data in a single model. This approach achieves state-of-the-art performance for Class I and Class II predictions.

## Materials and Methods

### Dataset

In order to train the Class I network, we pulled linear epitopes from the Immune Epitope Database (www.iedb.org) who had Class I restriction in humans with quantitative measurements of ic50 by purified MHC competitive radioactive and purified MHC competitive fluorescence assays, defining binding as peptide/allele pairings with ic50’s < 500nm. We transform the ic50 values by the equation (1) to scale from 0 to 1 where values below 1 nM are set to 1 nM and values above 50,000 nM are set to 50,000 nM.

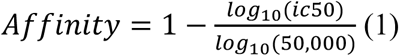

We additionally added another large data set of Class I binding predictions from *Kim et. al*(Kim *et al.*, 2014). We then restricted our training to entries where the full allele was provided (i.e. HLA-A*02:01) and then aggregated multiple peptide/allele pairings, taking the median value as a consensus where there were multiple peptide/allele pairings. In order to train the Class II network, we used a large data set published by *Jensen et. al*, following the same data preprocessing as described above(Jensen *et al.*, 2018).

For the purpose of comparing against other algorithms, we collected all the benchmarks from the IEDB automated server benchmarks for both Class I and Class II and restricted our analysis to benchmarks collected from competitive quantitative assays given our network was trained on data from these types of assays.

### Network architecture

The conventional neural network architecture as initially conceived for image classification tasks generally follows the format of stacking multiple convolutional layers with some type of non-linear activation and generally a max pooling step (Krizhevsky *et al.*, 2012). This approach transforms a photograph that is wide and tall with few features (RGB channels) to one that is compressed but with many features. The max pooling operations reduce the size of the photograph as it passes through the network but still maintain local spatial information. Applying this architecture to biological sequence analysis where peptides have variable lengths becomes problematic since neural networks require fixed size inputs. In the image classification world, this can be solved by rescaling or padding photographs so they all have the same pixel-by-pixel dimensions. Rescaling works well since RGB channels are continuous variables where down-sampling or interpolation algorithms can be applied but fails to translate to sequence analysis as sequences do not have a continuous numerical representation. In order to tackle this problem, *Vang et. al.* took an approach where they trained the network for a fixed size input. However, this approach would prevent training an entire allele’s set of peptides together which should significantly improve learning since the features between 9 and 10mers are most likely highly conserved. In order to be able to train a single model for an entire class of HLA molecules, we employed an approach where each peptide zero-padded (right) into a 15-mer window for Class I and 40-mer window for Class II. **(Figure 1A)**. This allows the network to take in sequences up to 15-mer in length for Class I predictions and up to 40-mer in length for Class II.

Since certain amino acids may share similar functional properties with others, we wanted to train an embedding that captures properties of each amino acid as the network is trained for prediction. In previous work, *Vang et. al* trained an embedding by using the Word2Vec algorithm to vectorize each amino acid based on its contextual use within the epitome. We chose to instead train the embedding with the classification task in mind as we believe this should learn the most salient embedding for the task at hand and our integrated approach would allow for the most amount of data to be leveraged towards training this embedding matrix.

In order to analyze this type of input to the network, we chose to use parallel convolutions of kernel length of 10-mers, knowing this should be large enough to encompass the 9-mer core that represents the length of the interacting peptide with the MHC molecule (Matsumura *et al.*, 1992). In comparison to other methods of biological sequence comparisons that use sequence alignment algorithms to assess conserved motifs, this network learns 1024 10-mer ‘trainable’ motif detectors (Sidhom *et al.*, 2017). The critical piece of the algorithm at this point is how it handles the max pooling step following these convolutions. Since our inputs can have variable lengths with zero-padding, if we chose a max-pooling strategy that divided the sequence into segments, we could be comparing segments in some windows where there was no sequence information to windows where there was sequence information **(Figure 1B)**. This would be especially problematic with Class II molecules where the input length can be highly variable. However, by conducting a global max-pooling operation, one is able to detect the relevant binding frame regardless of where it lies within the larger sequence.

**Figure 1:**
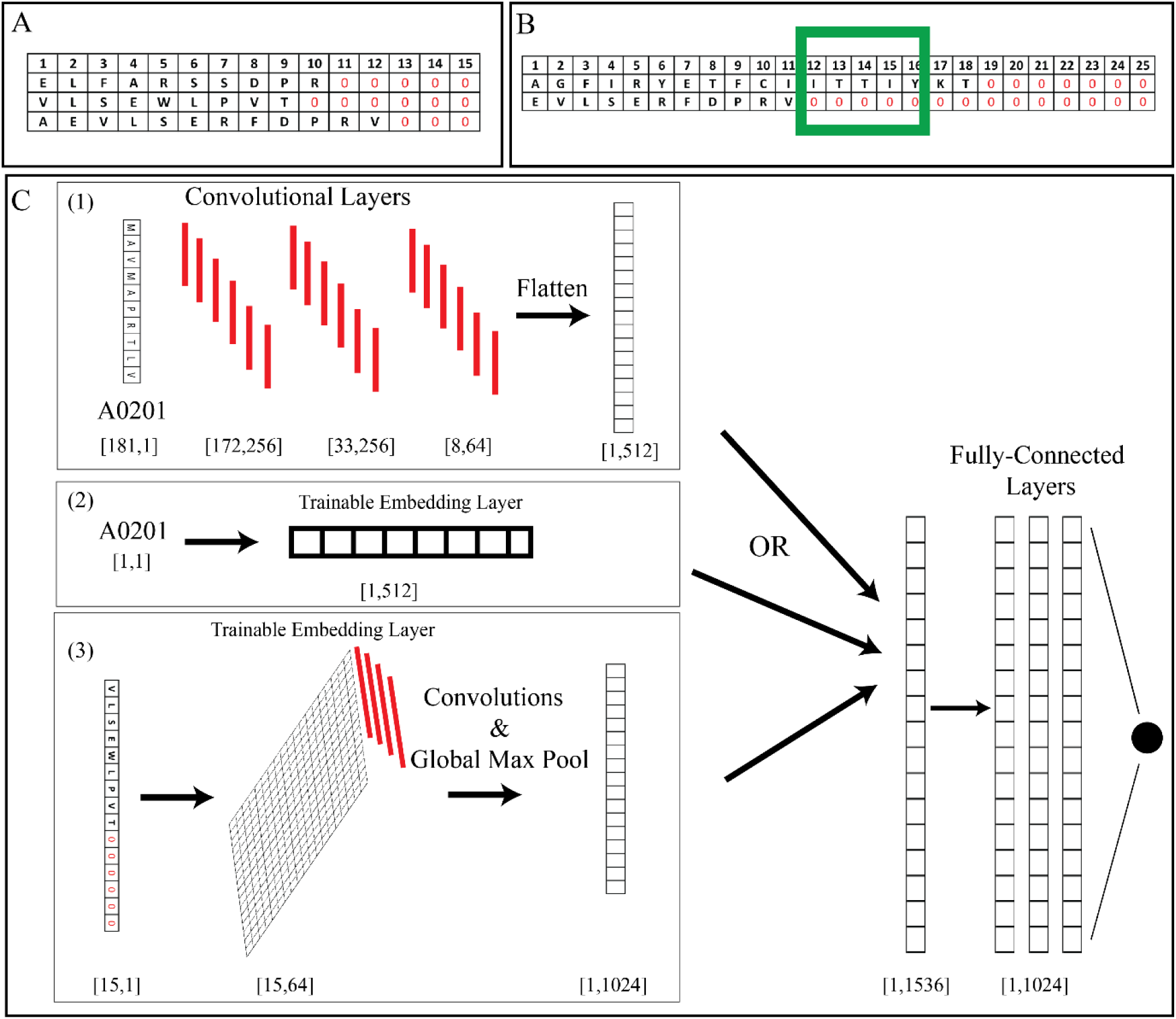
AI-MHC Architecture. A) Zero-padding scheme for handling variable length sequences. B) Green window highlights problem with max-pooling in segments as areas of null sequence can be directly compared to areas of real sequence. C) AI-MHC is designed to take a peptide/allele pair which are transformed with either (1) convolutional layers or (2) trainable embedding layers learning vector representations of alleles and (3) amino acids. 1024 10-mer convolutions with global max pooling are applied to the sequence resulting in a [1,1024] feature map for each sequence. The sequence feature map is then concatenated to the [1,512] allele feature map. This long-form vector [1,1536] is then followed by 3 fully-connected layers with 50% dropout and utilizing leaky relu activations functions with a final output node with sigmoid activation.

In order to train one unified architecture to predict binding based on sequence and allele input, we required an input to the network to be an allele paired to a given peptide. In order to integrate information about the allele into the network, we experimented with two methods: 1) applying convolutional layers to the actual protein sequences (Szolek *et al.*, 2014) of the MHC alleles to extract structural features of each allele **(Figure 1C-1)** or 2) training an embedding layer of 512 dimensions in order to learn properties of each allele **(Figure 1C-2)**. This is particularly advantageous because by training on a large dataset of all epitopes for various alleles, the net is able to learn features or train an embedding that can understand which alleles share similar properties and therefore, may share similar binding characteristics. Following either this convolutional feature extraction or embedding, this 512-dimensional vector is then joined with the 1024-dimensional feature vector for the sequence. In experimenting with both approaches, we found no difference in overall performance of the classifier and chose to implement a trainable embedding layer, as this was more computationally efficient. At this point, 3 fully connected layers (combining features extracted from MHC allele and peptide sequence) are implemented in which has final layer has a single output from a sigmoid activation, modeling the nM binding of the given MHC allele to peptide sequence pairing. The entire architecture was implemented with Google’s TensorFlow™ deep learning library.

## Results

### Neural Network Characterization

The presented neural network architecture contains two critical features that facilitate the use of the largest combined dataset of a given class of MHC for training; 1) translational invariance by convolutional layers that utilize a global max-pooling operation and kernel size that encompasses the entire possible length of the binding interaction between the peptide and MHC molecule and 2) integration of MHC allele as paired input with peptide sequence via an embedding layer, thereby enabling the entirety of the MHC Class I data to be used ensemble for training as compared to stratification by MHC allele. Besides creating larger datasets by combining data from different alleles and varying sequence lengths, the architecture developed allows the network to learn the properties of both amino acids and MHC molecules during its training, since each have a trained embedding matrix based on the data. In particular, an MHC embedding layer learns features of the MHC, allowing the model to learn from a larger dataset and translate knowledge of binding between alleles in the same supertype, sharing similar binding properties. To prove these points, we conducted two experiments that asses the invariance of the network as well as assess the quality of the embedding in its ability to cluster similar amino acids and HLA alleles and translate knowledge across alleles in the same supertype.

In order to test the invariance of the network, we created a synthetic dataset of 10,000 peptides of varying lengths between 8-11 amino acids resembling either A0201 binding peptides, L at P2, V at P9 (Drijfhout *et al.*, 1995), or a scrambled sequence containing a L and V at random positions **(Figure 2A)**. We provided the network the ability to learn one 10-mer feature, as this is the extent of information the network should need to make this classification correctly and were able to show that despite the L-V motif being placed in various frames in variable length peptides, our network was able to achieve perfect classification accuracy, as would be expected from a detect system that exhibited translational invariance **(Figure 2B**).

**Figure 2:**
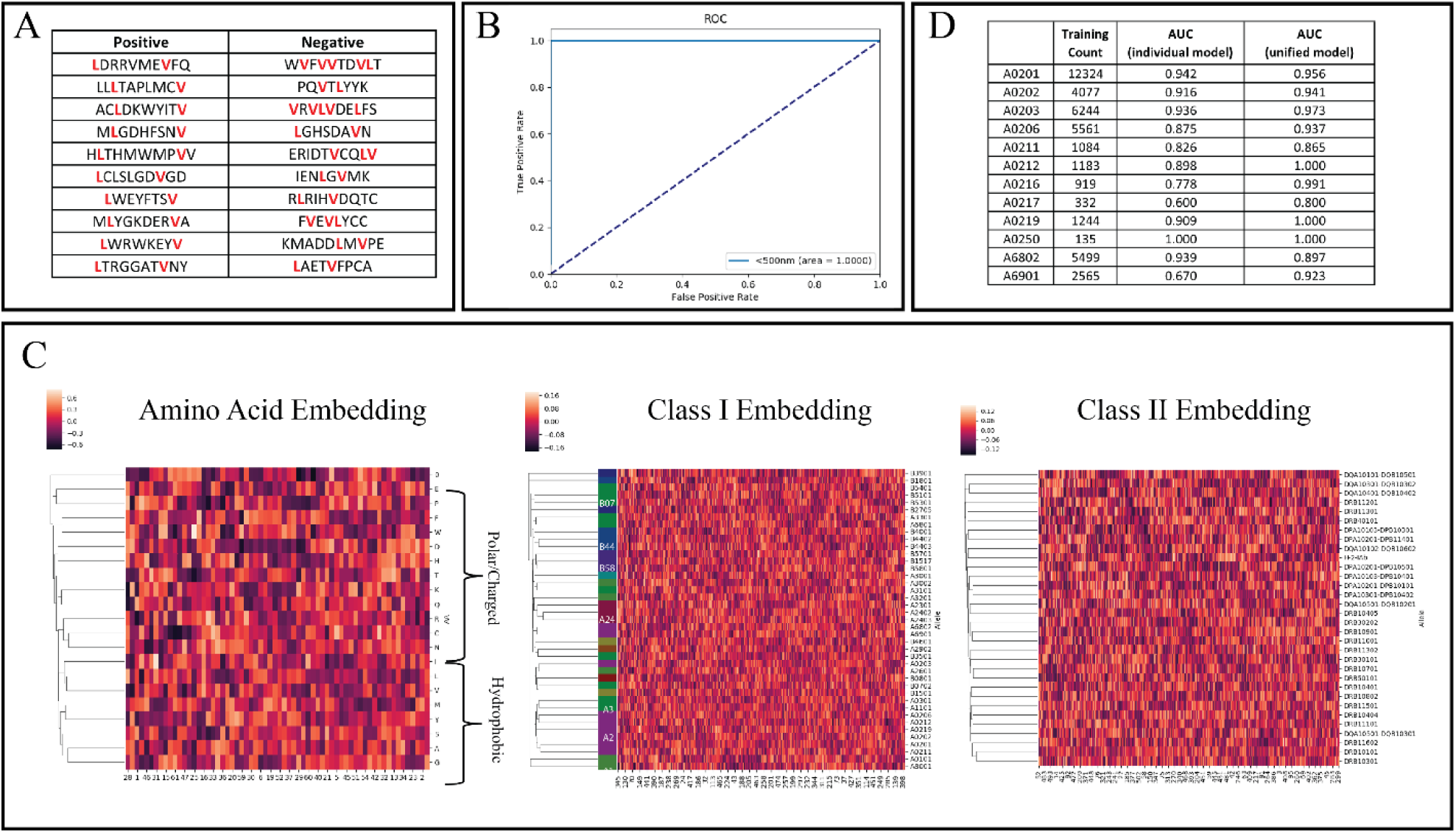
Network Characterization. A) Examples from synthetic dataset meant to mimic A0201 binding motifs (Leucine at P2, Valine at P9) in various frames within 8-11mer sequences. Red amino acids correspond to leucine and valine placed in correct and incorrect frames. B) Receiver Operating Characteristic of AI-MHC on synthetic dataset. C) Trained embedding layers were extracted from the network graph for amino acid, Class I, Class II embeddings and are visualized with clustermaps.

After training on MHC Class I and Class II data, we examined the embedding layers for both the amino acids and MHC alleles and noted that indeed amino acids with similar biophysical properties and MHC alleles in the same supertype **(Supplementary Table 1)** (Sidney *et al.*, 2008) were indeed clustered together **(Figure 2C)**, suggesting the network had learned which amino acids and HLA molecules share similar binding properties. While there does not exist a formal definition of supertypes for Class II molecules, we saw a similar clustering of related Class II molecules as well **(Figure 2C).** In order to assess the benefit of training an integrated model, we trained each of the sub-alleles of the HLA-A2 supertype in either individual or a unified model and compared their AUC values. We noted significant improvements in performance 10 of the 12 A2 supertype alleles **(Figure 2D)**, suggesting the network was able to translate its knowledge about the MHC-A2 supertype across its sub-alleles.

**Table 1:** AUC Values for Class I Alleles.

| Allele | Peptide Counts | AUC |
| --- | --- | --- |
| A0101 | 4609 | 0.950 |
| A0201 | 12324 | 0.956 |
| A0202 | 4077 | 0.941 |
| A0203 | 6244 | 0.973 |
| A0206 | 5561 | 0.937 |
| A0211 | 1084 | 0.865 |
| A0212 | 1183 | 1.000 |
| A0216 | 919 | 0.991 |
| A0217 | 332 | 0.800 |
| A0219 | 1244 | 1.000 |
| A0250 | 135 | 1.000 |
| A0301 | 7195 | 0.968 |
| A1101 | 6248 | 0.970 |
| A2301 | 2416 | 0.953 |
| A2402 | 3191 | 0.995 |
| A2403 | 1227 | 0.990 |
| A2501 | 960 | 1.000 |
| A2601 | 4307 | 0.956 |
| A2602 | 631 | 0.955 |
| A2603 | 522 | 1.000 |
| A2902 | 2548 | 0.980 |
| A3001 | 2717 | 0.857 |
| A3002 | 1847 | 0.775 |
| A3101 | 5621 | 0.968 |
| A3201 | 1089 | 1.000 |
| A3215 | 74 | 1.000 |
| A3301 | 3510 | 0.963 |
| A6601 | 52 | 1.000 |
| A6801 | 3708 | 0.960 |
| A6802 | 5499 | 0.897 |
| A6901 | 2565 | 0.923 |
| A8001 | 1164 | 1.000 |
| B0702 | 4513 | 0.937 |
| B0801 | 3298 | 0.924 |
| B0802 | 1000 | 0.983 |
| B1501 | 4178 | 0.963 |
| B1503 | 594 | 0.814 |
| B1517 | 1446 | 0.945 |
| B1801 | 2594 | 0.896 |
| B2705 | 3433 | 0.897 |
| B3501 | 3198 | 0.982 |
| B3801 | 492 | 1.000 |
| B3901 | 1623 | 0.906 |
| B4001 | 3199 | 0.992 |
| B4002 | 964 | 0.933 |
| B4402 | 2117 | 0.991 |
| B4403 | 1382 | 1.000 |
| B4501 | 953 | 0.909 |
| B4601 | 1806 | 0.955 |
| B5101 | 2718 | 0.956 |
| B5301 | 1616 | 0.938 |
| B5401 | 1110 | 1.000 |
| B5701 | 2781 | 0.963 |
| B5801 | 3118 | 0.978 |
| B8301 | 333 | 1.000 |

### Class I Metrics

We collected a total of 148,540 unique allele/peptide pairing with ic50 values from the IEDB (www.iedb.org) and a previously published dataset by *Kim et.al* spanning 86 HLA-A,B,C,E alleles **(Supplementary Table 2)**^*i*^. For the purpose of training, we split these data sets into a train set of 95% and split the remaining 5% for validation and testing, resulting in a train size of 141,113 peptides, a validation size of 3,713, and a test size of 3,714. The network was trained on the train data while validation data was used to determine when to stop training the neural network. In this set of peptides, our model achieved an overall AUC of 0.956 on our internal independent test set **(Figure 3A)**, generally achieving higher AUC values for where there was more data available for a given MHC molecule **(Table 1)**.

**Table 2:**
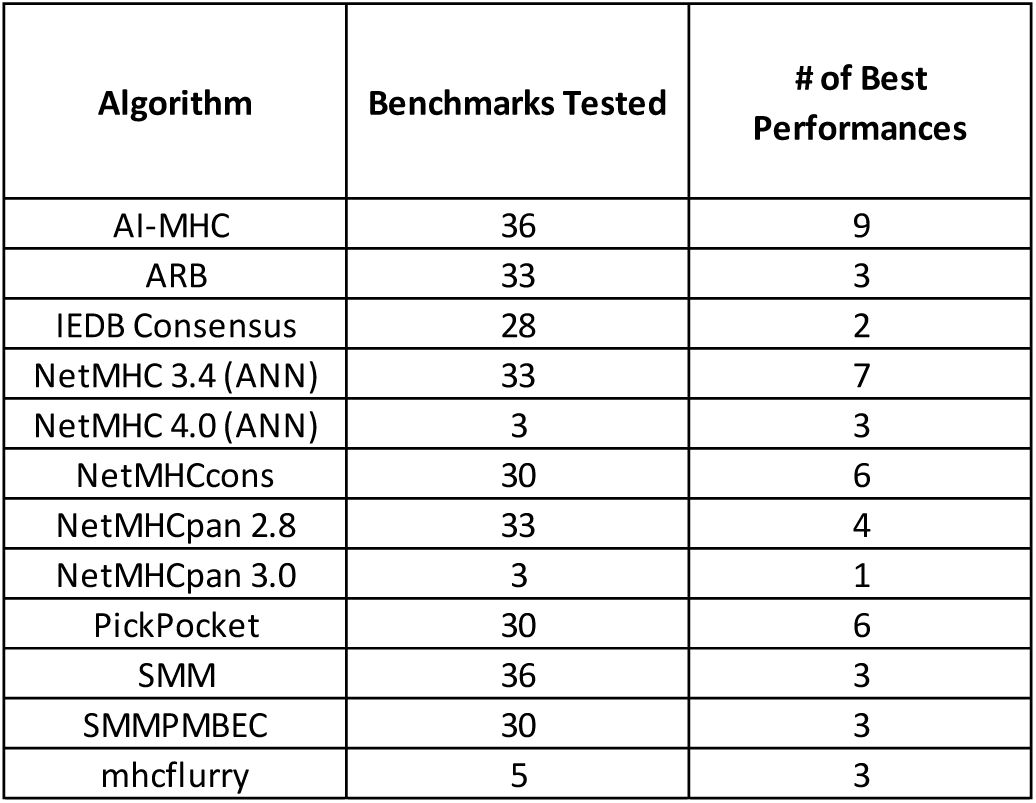
Class I – IEDB Benchmark Performance. We collected all benchmark datasets from the IEDB for which our algorithm had at least 10 training examples for the allele tested to assess performance against 11 of the available algorithms. # of Best Performances refers to the number of benchmarks a given algorithm ‘won’.

| Algorithm | Benchmarks Tested | # of Best Performances |
| --- | --- | --- |
| AI-MHC | 36 | 9 |
| ARB | 33 | 3 |
| IEDB Consensus | 28 | 2 |
| NetMHC 3.4 (ANN) | 33 | 7 |
| NetMHC 4.0 (ANN) | 3 | 3 |
| NetMHCcons | 30 | 6 |
| NetMHCpan 2.8 | 33 | 4 |
| NetMHCpan 3.0 | 3 | 1 |
| PickPocket | 30 | 6 |
| SMM | 36 | 3 |
| SMMPMBEC | 30 | 3 |
| mhcflurry | 5 | 3 |

**Figure 3:**
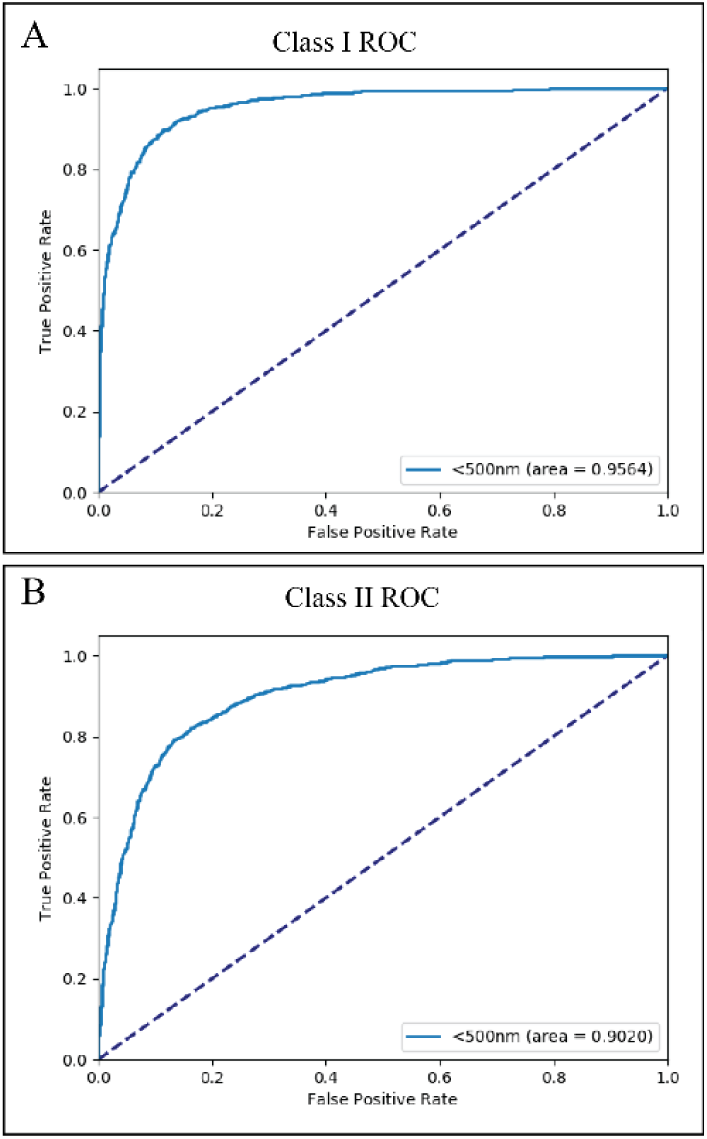
ROC for Class I and Class II models.

In order to gauge where our algorithm stood against the current state-of-the-art algorithms, we pulled all the ic50 benchmarks from the IEDB (www.tools.iedb.auto_bench/mhci/weekly/) to assess the performance of our algorithms against the 11 provided algorithms. Since it is unclear whether these are considered independent benchmarks as the IEDB cannot verify that the tested allele/peptide pairings have not been seen by the benchmarked algorithms, it is difficult to truly compare performance at an algorithmic level. Nonetheless, we removed all records in the benchmarks from our training data before assessing the performance of our models. Of the 47 available benchmarks, AI-MHC performed the best on 9/36 datasets (next highest was NetMHC 3.4 with 7/36) where we had at least 10 peptide examples for training to a given MHC allele. **(Table 2)**. Full comparison of all algorithms on all benchmarks in **Supplemental Table 3.**

**Table 3:**
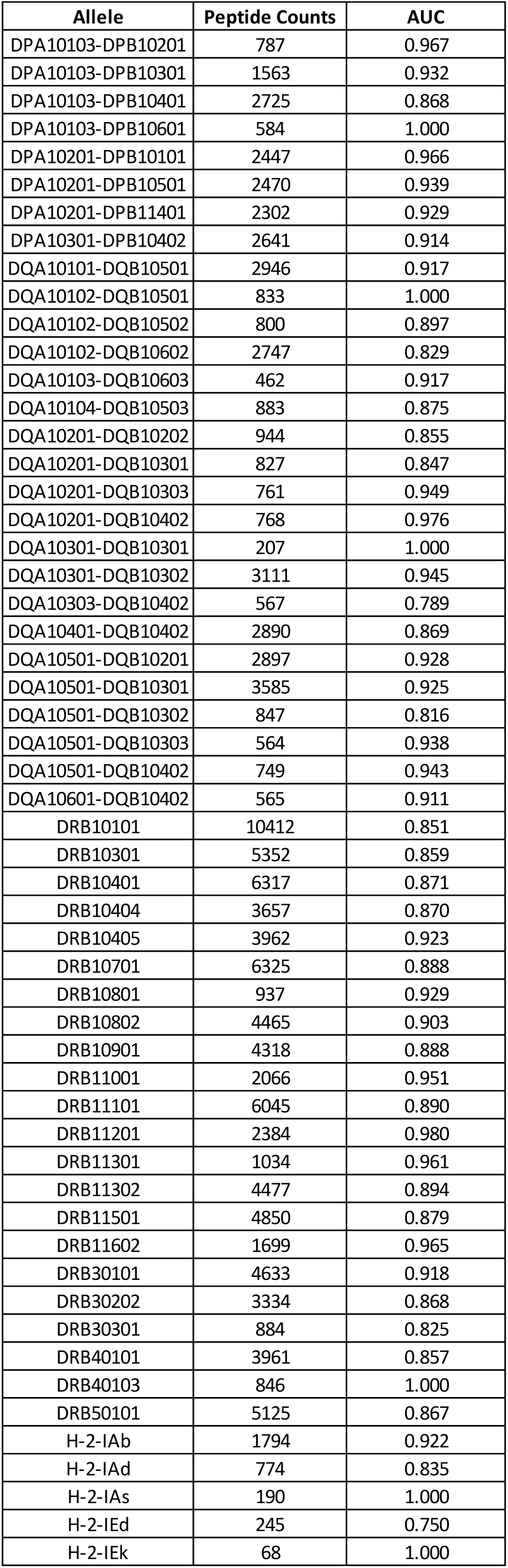
AUC Values for Class II Alleles.

### Class II Metrics

In order to test whether this type of architecture would also be relevant in predicting Class II binding, we collected a total of 134,281 unique allele/peptide pairings with ic50 values from a previously published dataset by *Jensen et.al* spanning 80 alleles **(Supplementary Table 4)**. For the purpose of training, we again split this data set the same way as with the Class I training, resulting in a train size of 127,566 records, a validation size of 3,357, and a test size of 3,358. Training was completed in the same way as described above. In our internal independent data set, our model achieved an overall AUC of 0.902 **(Figure 3B & Table 3)**. In comparison to AUC values published by *Jensen et.al* on the same data set, our model outperforms all recorded AUC values from the NetMHCIIpan-3.2 (AUC = 0.858, 0.861, 0.826). Furthermore, we pulled all ic50 benchmarks from the IEDB (http://tools.iedb.org/auto_bench/mhcii/weekly/) to assess the performance of our algorithm against the 6 provided algorithms. Once again, we benchmarked our algorithm by removing entries from our training set that appeared within the IEDB benchmarks. Of the 54 available benchmarks, our model performed the best on 18/48 datasets which we had at least 10 training examples, the second highest number of best performances to NetMHCIIpan-3.1 **(Table 4)**. Full comparison of all algorithms on all benchmarks in **Supplemental Table 5.**

**Table 4:** Class II – IEDB Benchmark Performance. We collected all benchmark datasets from the IEDB for which our algorithm had at least 10 training examples for the allele tested to assess performance against 11 of the available algorithms. # of Best Per

| Algorithm | Benchmarks Tested | # of Best Performances |
| --- | --- | --- |
| AI-MHC | 48 | 18 |
| Comblib matrices | 18 | 0 |
| Consensus IEDB method | 36 | 6 |
| NN-align | 34 | 2 |
| NetMHCIIpan-3.1 | 48 | 22 |
| SMM-align | 34 | 3 |
| Tepitope (Sturniolo) | 29 | 1 |

## Conclusion

In this work, we present an integrated deep learning architecture to predict MHC Class I and Class II binding, able to achieve state-of-the-art performance through utilizing innovative changes in architecture allowing the network to be trained on effectively larger datasets, which is a well-known requirement to better training deep neural networks. This is accomplished through training an entire class of MHC alleles in a unified model by learning an embedding layer for the allele allowing leveraging of binding information between alleles of the same supertype. Furthermore, the architecture becomes flexible to sequence input length by utilizing a global max-pooling operation across the input peptide sequence following convolutions to achieve translational invariance where a frame of interest needs to be learned in the context of a longer, variable length peptide sequence.

In attempting to assess the performance of our algorithm, we noted the difficulty in making equivalent comparisons to other algorithms in the field as there are no clear train/test datasets that all algorithms can be benchmarked against in sense that we could determine what data should be training versus independent test for any given algorithm. In an attempt to conduct a robust analysis, we first created an internal independent test set that was not used for training purposes, as per our methods section. Our results for Class I and Class II (AUC – Class I = 0.956 & AUC – Class II = 0.902) suggest our algorithm is one of the top performing algorithms. However, without the ability to train/test other algorithms, it is not possible to directly assess exactly where our algorithm stood. For Class I assessment, even when removing all IEDB examples from our training, our algorithm still had the highest number of best performances on benchmarks where we had sufficient training examples. For Class II, we felt we were able to make a fairer comparison as NetMHCIIpan-3.2 released a large dataset on which they trained/tested. By using the same data and their reported AUC values, we were confident that our algorithm was truly out-performing what is considered the best Class II prediction algorithm by ∼4% AUC despite not having more ‘best performances’ in the IEDB benchmarks. While our dilemma in assessing performance against other algorithms is not a new one, we suggest that there needs to be a method, such as the annual ImageNet Challenge, by which algorithms can be compared in a fair way where training/testing datasets are equivalent across all algorithms to truly assess the best algorithmic approaches. That being said, given the volume of data we collected for MHC class I and II in conjunction with the ability of our algorithm to more fully leverage each of the sets of data in total, our approach of isolating an internal independent test set still allows for the evaluation of performance across thousands of allele/peptides.

Finally, we have provided a user-friendly website for use of our algorithms for both Class I and Class II predictions with performance metrics provided for each allele. We believe this level of transparency in the allele-level performance is important to better inform the user of the confidence in any prediction based on the number of peptides tested and the internal AUC achieved for any given allele.

